# Fast and simple comparison of semi-structured data, with emphasis on electronic health records

**DOI:** 10.1101/293183

**Authors:** Max Robinson, Jennifer Hadlock, Jiyang Yu, Alireza Khatamian, Aleksandr Y. Aravkin, Eric W. Deutsch, Nathan D. Price, Sui Huang, Gustavo Glusman

## Abstract

We present a locality-sensitive hashing strategy for summarizing semi-structured data (e.g., in JSON or XML formats) into ‘data fingerprints’: highly compressed representations which cannot recreate details in the data, yet simplify and greatly accelerate the comparison and clustering of semi-structured data by preserving similarity relationships. Computation on data fingerprints is fast: in one example involving complex simulated medical records, the average time to encode one record was 0.53 seconds, and the average pairwise comparison time was 3.75 microseconds. Both processes are trivially parallelizable.

Applications include detection of duplicates, clustering and classification of semi-structured data, which support larger goals including summarizing large and complex data sets, quality assessment, and data mining. We illustrate use cases with three analyses of electronic health records (EHRs): (1) pairwise comparison of patient records, (2) analysis of cohort structure, and (3) evaluation of methods for generating simulated patient data.

## Background

Fast comparison of a large number of semi-structured data records can be valuable for analysis of any knowledge domain. Electronic healthcare records (EHRs) present an important real-world test case because many healthcare questions can benefit from finding the similarity between different records. Examples include finding candidates for clinical trials, evaluating treatment response, and investigating cost variation among patients with similar health situations. Currently, approaches to identifying similar records include (1) designing semantically-informed comparison on structured elements of interest, such as demographics, conditions or medications and (2) using semantically and structurally agnostic file comparison. The first requires explicit mapping of domain knowledge to data structure, and the second becomes resource intensive when scaled to millions of patient records. Here, we present a fast, semantically and structurally agnostic method for analyzing semi-structured data, and investigate its application to several basic analyses of patient records.

HL7 Fast Healthcare Interoperability Resources (FHIR) [1] is a structured data standard with well-defined schema. Other examples of structured data include tabular, relational and flat file formats that have a data dictionary. FHIR also supports semantic interoperability at Healthcare Information and Management Systems Society (HIMSS) Level 3 [2], using terminologies such as SNOMED CT [3]. FHIR can be represented in JavaScript Object Notation (JSON), a semi-structured format with a “self-describing”, flexible schema [4]. Other commonly-used flexible schemas include XML and ASN.1 [5]. FHIR and JSON can also include minimally structured data. For example, radiology impressions are free text, although they generally follow specific conventions and use a constrained subset of precise medical terms. Patient encounter notes may have almost no explicit structure, but they usually follow conventions that can guide domain-specific comparisons. Such minimally structured data could be converted into semi-structured data prior to downstream analysis.

We describe here a novel strategy for encoding semi-structured data into “data fingerprints” that facilitate their efficient comparison. This strategy has potential applications in many domains; we present here examples of its use for fast and simple comparison of EHRs.

## Results and Discussion

### Overview of the strategy

We have developed a strategy for compressing semi-structured data into a greatly reduced form which preserves similarity and is suitable for comparison of data objects, without preserving the ability to recreate the details or structure of the original data. This strategy proceeds in three stages. First, the semi-structured data object is parsed to identify hierarchical elements of local structure, corresponding to nested *objects* (key/value pairs), *arrays* (ordered lists of elements), and *values*. These elements are transformed into a series of *statements* (Figure 1) that describe the properties of the objects (e.g., “object X has a property Y with value Z”, where Z can be a simple value or a nested element) and the ordering of elements in the arrays (e.g., “in array X, element Y is followed by element Z”). Then, each statement is encoded into a *vector contribution*; this involves specific procedures for encoding different data types (e.g., numbers and strings) and for combining the three elements in a statement into the final vector contribution (see Methods). Finally, the vector contributions are summed to yield the *raw fingerprint*, which can then be centered and scaled to yield the *normalized fingerprint*.

**Figure 1.**
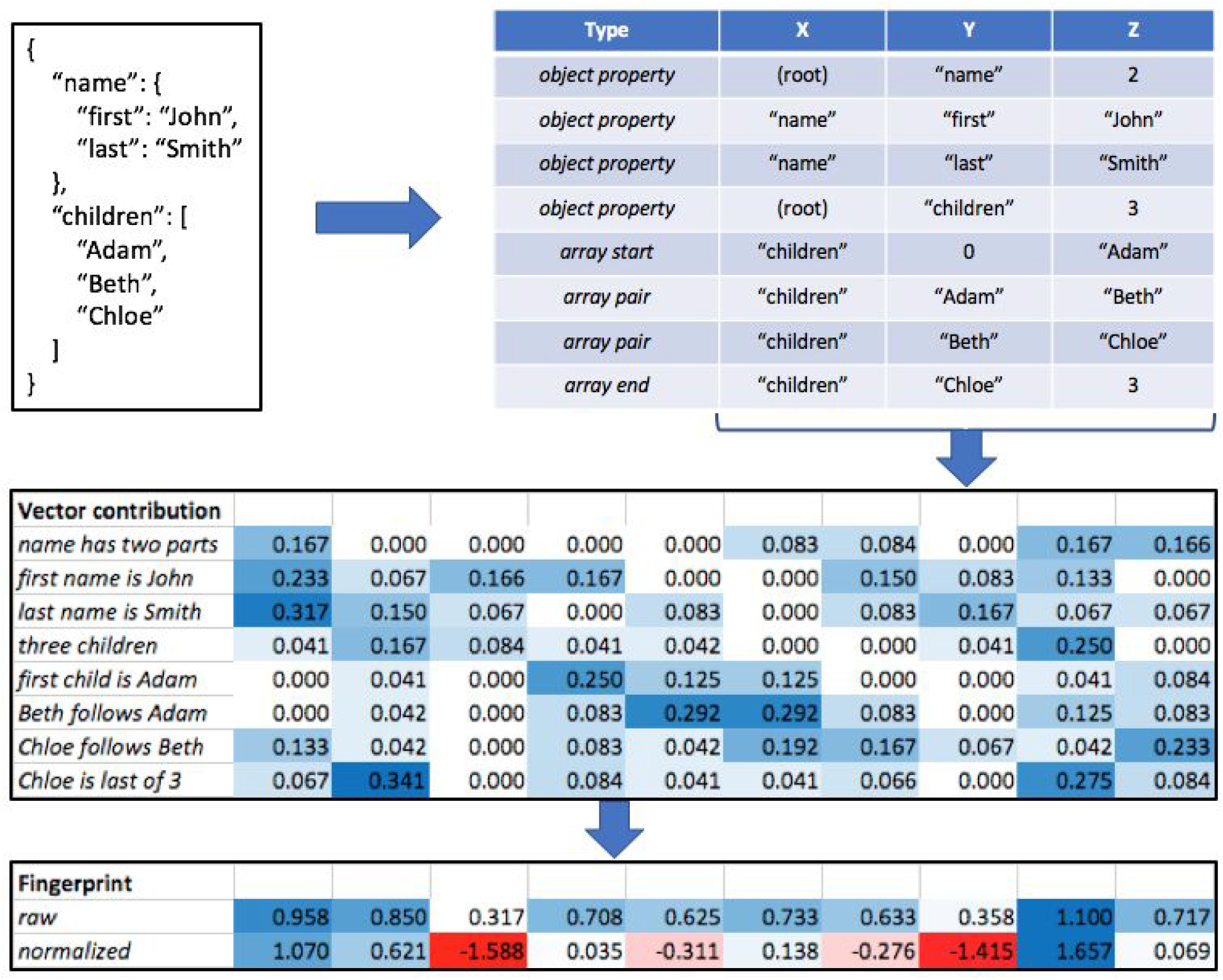
Overview of the strategy. Semi-structured data (here shown as a JSON object, upper left) is transformed into a series of statements (upper right) describing local substructure within objects and arrays. Each statement is then independently encoded into a numerical vector (in the example at the center, of length 10) as described in the Methods. The sum of these vector contributions (bottom) yields the ‘raw’ fingerprint, which can then be centered and rescaled to yield the ‘normalized’ fingerprint.

### Data fingerprints are suitable for studying complex medical records

SyntheticMass is an open-source, simulated Health Information Exchange populated with realistic “synthetic residents” of Massachusetts [6], with population health and demographic data at the city and town level, and fictional but realistic data representing synthetic patients. We studied a sample set of 116,185 simulated patient records from SyntheticMass in FHIR Draft Standard for Trial Use (DSTU) 3 format. We computed data fingerprints for each simulated patient (fingerprint length *L*=200); average fingerprint computation time was 0.53 seconds/record. We evaluated the structure of the remaining cohort of 116,185 patients using PCA on the normalized fingerprints (Figure 2).

**Figure 2.**
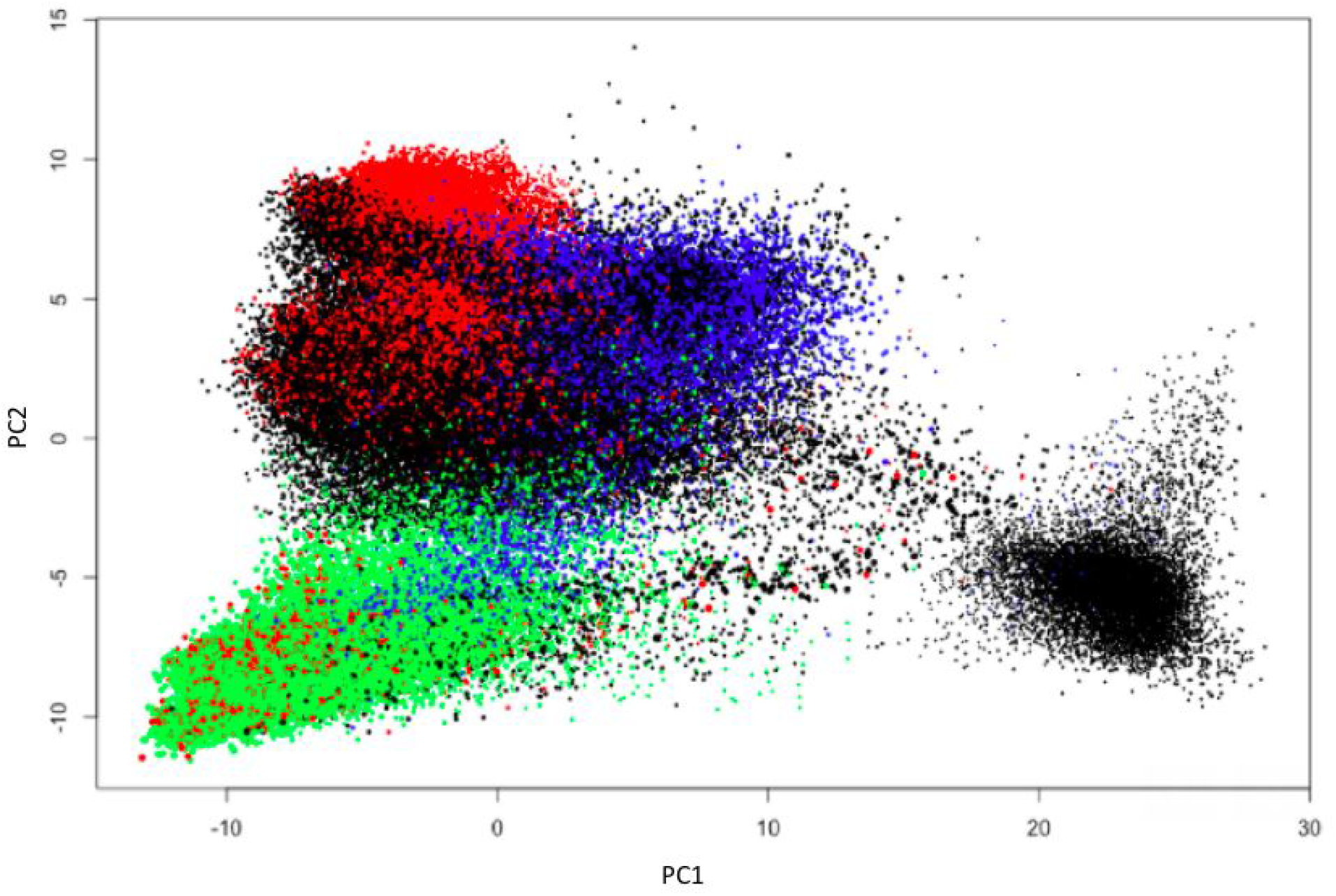
Observed structure in the SyntheticMass sample data set. Each circle represents one simulated patient; circle area is proportional to the number of statements in the patient’s data fingerprint. PC1 and PC2: first and second principal components, respectively. Colors denote presence of specific terms in the simulated patient records; blue: ‘Normal pregnancy’ or ‘Prenatal visit’; red: ‘DTaP’ or ‘Pneumococcal conjugate’; green: ‘Basic Metabolic Panel’; black: all other simulated patients. The separate cluster to the right includes most simulated patients with the term ‘Death Certification’.

We then contrasted the frequency of terms observed in simulated patient records across the different clusters derived from the PCA, and identified many terms that are enriched in one cluster relative to other clusters. We highlight a few of these in Figure 2. In particular, we observe clusters enriched in simulated patients with pregnancy-related terms, with vaccination-related terms, with information on standard metabolic panels, and (as a largely distinct cluster) with records on patients simulated to be deceased.

### Pairwise comparison of records via data fingerprints

We computed all possible pairwise correlations among a subsample of 1462 simulated patients in the SyntheticMass sample cohort (1,067,991 pairwise comparisons); this analysis took 4 CPU seconds, i.e., an average of 3.75 microseconds per comparison. All the pairwise correlations among patients in the cohort are very high (ranging from 0.842 to 0.997), most likely reflecting the nature of the JSON FHIR format. For example, every record is made up of resources, each of which has a JSON key for “resourceType”, and contains common data types, such as “dateTime”, or “coding”.

**Table 1.**
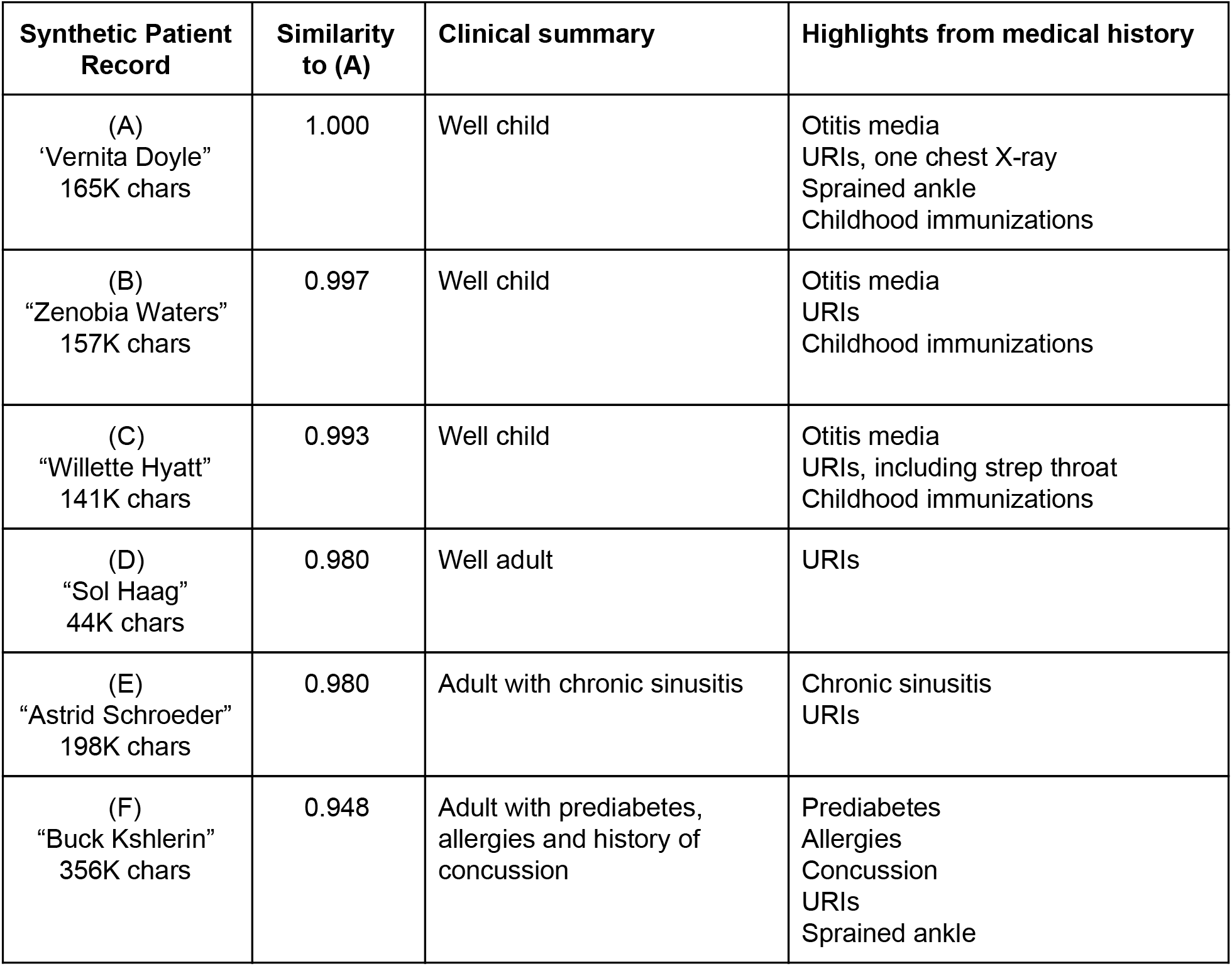
Pairwise similarity (fingerprint Spearman correlation) between synthetic Patient A and synthetic Patients B through F; their simulated names (in quotes) serve as identifiers in the SyntheticMass database. A and B are the closest pair from collection of 1462 records. Records are ordered by distance from Record A. Impression: Clinical impression from manual review of text and display fields from FHIR JSON records. EHRs. URIs = upper respiratory infections. Chars = characters in file, not including spaces.

| Synthetic Patient Record | Similarity to (A) | Clinical summary | Highlights from medical history |
| --- | --- | --- | --- |
| (A)<br>"Vernita Doyle"<br>165K chars | 1.000 | Well child | Otitis media<br>URIs, one chest X-ray<br>Sprained ankle<br>Childhood immunizations |
| (B)<br>"Zenobia Waters"<br>157K chars | 0.997 | Well child | Otitis media<br>URIs<br>Childhood immunizations |
| (C)<br>"Willette Hyatt"<br>141K chars | 0.993 | Well child | Otitis media<br>URIs, including strep throat<br>Childhood immunizations |
| (D)<br>"Sol Haag"<br>44K chars | 0.980 | Well adult | URIs |
| (E)<br>"Astrid Schroeder"<br>198K chars | 0.980 | Adult with chronic sinusitis | Chronic sinusitis<br>URIs |
| (F)<br>"Buck Kshlerin"<br>356K chars | 0.948 | Adult with prediabetes,<br>allergies and history of<br>concussion | Prediabetes<br>Allergies<br>Concussion<br>URIs<br>Sprained ankle |

The most highly correlated pair of records involved simulated patients “Vernita Doyle” and “Zenobia Waters”. Table 1 lists the similarities and clinical summaries for these two, and other representative simulated individuals. We observe that the decreasing levels of correlation match the subjective intuition of similarity between patients. Of note, a single patient condition can have a significant impact on the content of a record. For example, Patient F had a FHIR Condition resources for Allergies, and other related content: allergy screening test, allergic disorder initial assessment, allergen immunotherapy drugs, an epinephrine auto-injector, and associated billing codes. This suggests that even if two patient records with a clinically similar conditions used a different SNOMED code in the FHIR Condition field, the similarity between the two records might still be detected by other elements within the larger signature for the condition, such as diagnostic maneuvers, therapy, and billing.

Detecting clinical similarity in real-world records that were created in different healthcare systems will likely be more difficult than in the small set of synthetic data records we evaluated here. Additionally, the manual review we performed focused on whether patient records appeared to be similar from a clinical perspective. Since the method is domain-agnostic, the computed similarities could also reflect on other aspects of healthcare. For example, patients may have similar clinical situations but different healthcare encounter types and billing (ambulatory care vs emergency visit). Alternately one may identify individual records that show evidence of billing errors, and then find similar records that may also need correction. It will also be valuable to assess our strategy on records that include free text, such as encounter notes and diagnostic reports.

### Data fingerprints reveal cohort structure

The St. Jude Lifetime Cohort (SJLIFE) [7] is an unparalleled resource to longitudinally track health outcomes among childhood cancer survivors and community controls that is supported by a robust institutionally-funded clinical and research infrastructure at St. Jude Children’s Research Hospital. SJLIFE cohort has characterized over 3,000 pediatric cancer survivors by careful clinical ascertainment of phenotypes and whole genome and whole exome sequencing data. It is by far the single largest medically evaluated cohort of childhood cancer survivors in the world. The well-characterized clinical data in SJLIFE cohort covers all pediatric cancer subtypes [8], recruitment of participants representing the spectrum of pediatric, adolescent and young adult cancers as well as frequency-matched community controls, collection of comprehensive treatment data on all participants, provision of protocol-based medical assessments, assessment of patient-reported outcomes, validation of self-reported medical events, performance of periodic longitudinal evaluations, and collection of biologic specimens. Longitudinal systematic medical assessment of this unique cohort will facilitate elucidation of the pathophysiology of cancer treatment-related morbidity, identification of biomarkers of subclinical organ dysfunction, and characterization of high-risk groups who may benefit from interventions to preserve health.

We used data fingerprints to study electronic health records for 500 patients from the SJLIFE cohort (SJL500). These records, which are smaller and less complex than those in the SyntheticMass cohort, include seven main rubrics:

1. Population overview: sex, race, education, income, age, height, weight, age at diagnosis, smoker status, etc.
2. Chronic conditions: the maximal severity grade for each chronic condition evaluated for each individual.
3. Chemotherapy overview: for each of 104 chemotherapy agents, whether the patient received it within five years of primary diagnosis, or within ten years, or at any stage.
4. Chemotherapy dosage: for each chemotherapy agent, the dose received within five years of primary diagnosis, or within ten years, or at any stage.
5. Dosimetry: the maximum treatment dose to any portion of a body region (brain, neck, chest, abdomen, pelvis, total body irradiation, traditional mantle field, etc.
6. Radiation overview: administered radiation therapy potentially impacted body region (abdomen, brain, breast, chest, ear, eye, female genitals, gastrointestinal/hepatic, lung, male genitals, muscle, neck/thyroid, oral, pelvis, urinary).
7. Secondary primary cancer: details on any observed secondary cancers, not metastases of the initially diagnosed cancer.

**Figure 3.**
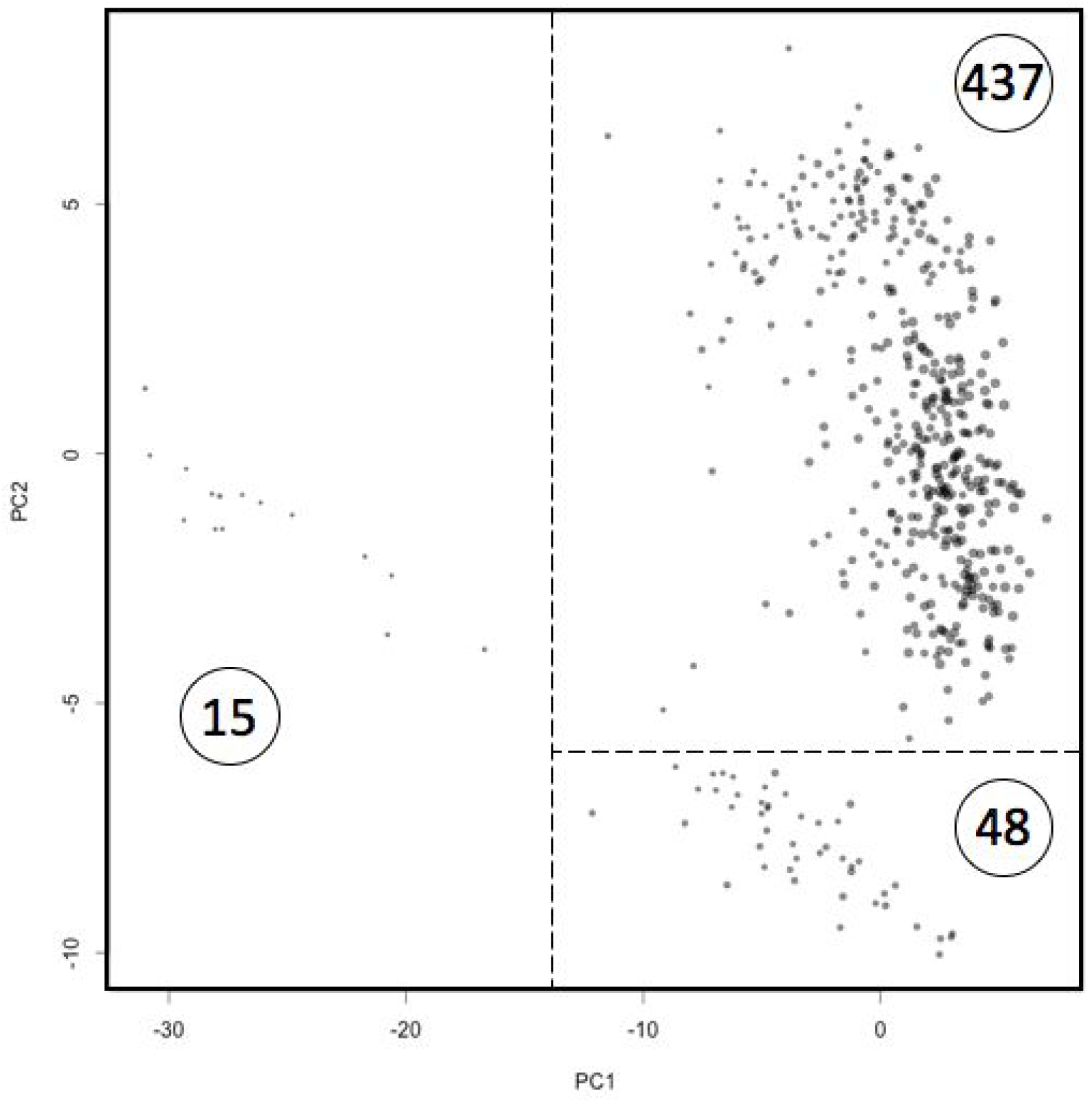
Structure in the SJL500 cohort. The data fingerprints for the 500 patients are separated into three clusters by Principal Component Analysis. Each circle represents one patient; circle area is proportional to the number of statements in the patient’s data fingerprint. PC1 and PC2: first and second principal components, respectively. Dashed lines represent the cutoffs used to define the clusters.

We computed data fingerprints for each SJL500 patient (fingerprint length *L*=200); average fingerprint computation time was 0.017 seconds/record. We evaluated the structure of the cohort using Principal Component Analysis (PCA) on the normalized fingerprints. This analysis revealed the presence of three main clusters in the SJL500 cohort (Figure 3). The first principal component (PC1) separates 15 patients ("small cluster") for which there is relatively little information in the data set. The second principal component (PC2) then separates between a set of 48 patients ("medium cluster") and the rest of the cohort ("large cluster" of 437 patients, with some further internal structure not discussed here).

We then contrasted the medium cluster and the large cluster by term enrichment analysis. We observed that several terms are much more frequently observed in the medium cluster than in the large cluster (Table 2).

**Table 2.** Enrichment of clinical terms in the SJL500 cohort when comparing the medium cluster (N=48) with the large cluster (N=437).

| Term | medium (N=48) | large (N=437) | Enrichment |
| --- | --- | --- | --- |
| chronic.Endocrine_Adrenalinsufficiency | 25.0% | 5.0% | 5.0x |
| chronic.Neurology_Cerebellardysfunction | 14.6% | 3.0% | 4.9x |
| chronic.Endocrine_ChildhoodGHD | 27.1% | 5.9% | 4.6x |
| chronic.Gastrointestinal_Esophagealstricture | 16.7% | 4.6% | 3.6x |
| chronic.Cardiovascular_VascularDisease | 16.7% | 5.5% | 3.0x |
| chronic.Neurology_Cranialnervedisorder | 10.4% | 3.4% | 3.0x |
| chronic.Pulmonary_Obstructivesleepapnea | 12.5% | 4.8% | 2.6x |
| chronic.Reproductive_Centralhypogonadism | 16.7% | 6.4% | 2.6x |
| chronic.Endocrine_Hypothyroidism | 66.7% | 27.0% | 2.5x |
| chronic.Neurology_Cerebrovascularaccident | 14.6% | 5.7% | 2.5x |
| chronic.Endocrine_Thyroidcyst/nodule | 29.2% | 13.5% | 2.2x |
| chronic.Endocrine_AdultGHD | 31.2% | 16.7% | 1.9x |
| radyn.potential_oral_10 | 91.7% | 49.0% | 1.9x |
| radyn.potential_oral_prim | 91.7% | 49.2% | 1.9x |
| dosimetry.mdart_neck_5 | 58.3% | 32.0% | 1.8x |
| dosimetry.mdart_neck_5_dose | 58.3% | 32.0% | 1.8x |
| radyn.potential_oral | 91.7% | 49.9% | 1.8x |
| radyn.potential_oral_5 | 89.6% | 48.7% | 1.8x |

### Data fingerprints facilitate evaluation of methods for generating synthetic data

Synthetic data generation (SDG) methods have been evaluated in multiple ways, ranging from simple statistical analysis of data size and distribution of values, to more complex considerations of realism and suitability for the purpose for which the simulated records will be used [6]. We previously published a method for generating realistic artificial genomic sequences [9] and their evaluation using a collection of descriptors (composition and complexity measures). We show here a similar application of data fingerprints to evaluate SDG methods in terms of the overall structure of the cohorts they produce.

We applied three methods for generating artificial records based on the real data from the SJL500 cohort (see Methods). The Random method samples from the observed values in the cohort but does not preserve the pattern of sparseness of each individual record. The Patterned method preserves the pattern of observed values for each individual, and randomizes the actual values observed. The Frequent method also preserves the pattern of observations for each individual, but replaces the observed value for each parameter with the most commonly observed value for that parameter. We computed data fingerprints (*L*=200) for all real and simulated records and compared them using PCA (Figure 4). As expected, the Random method yields very similar records that do not reconstruct the original cohort structure. The Patterned and Frequent methods yield cohorts that much more closely mimic that of the real data. Nevertheless, the cohort produced by the Frequent method is clearly distinguished from the other cohorts by a higher principal component (PC4), suggesting this method would be unsuitable as a substitute for the real data for downstream analytic applications. Thus, data fingerprints provide a simple and convenient method for evaluating the structure of cohorts produced by various simulation methods, and enable their comparison to the real data.

**Figure 4.**
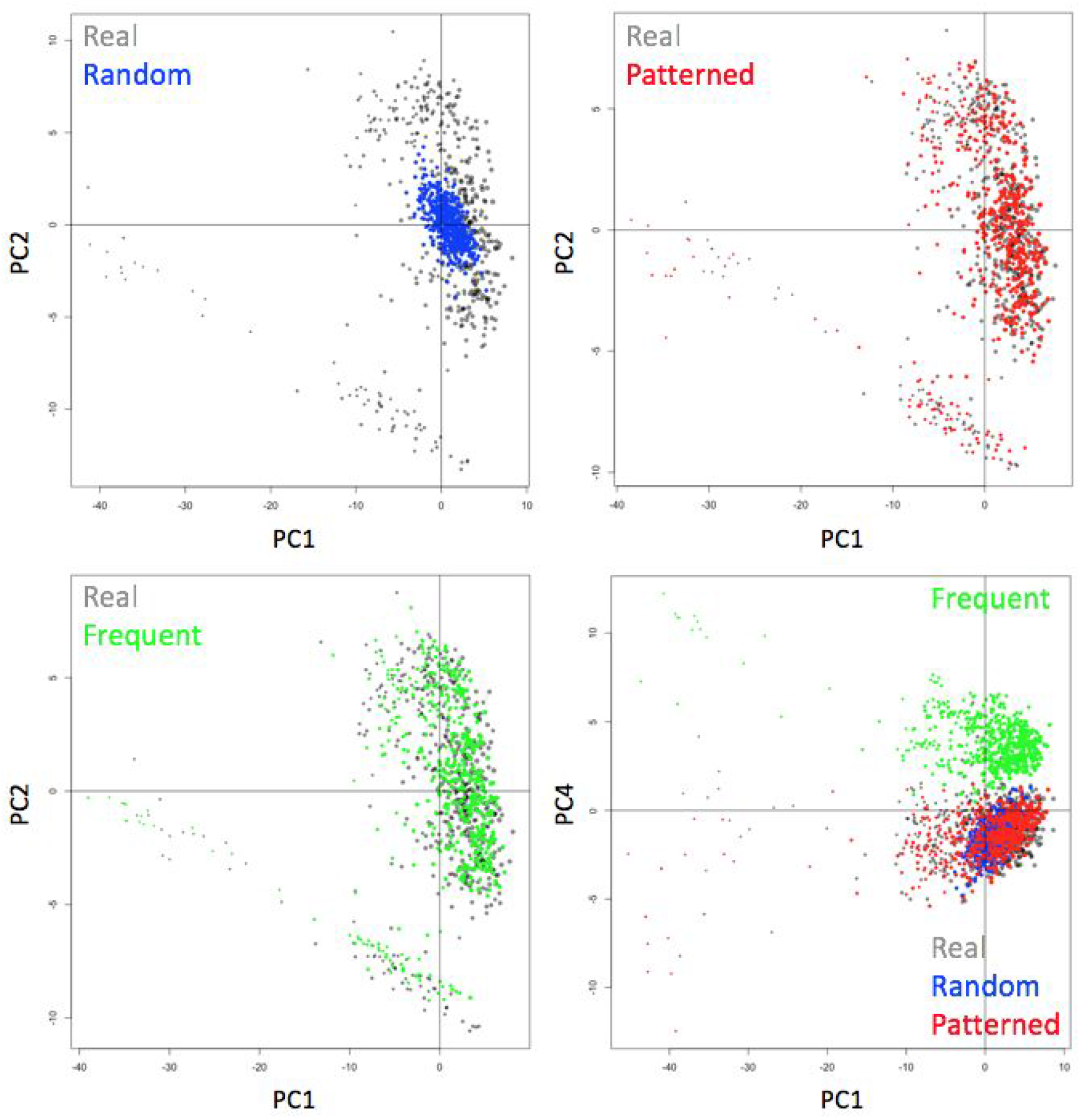
Comparison of artificial data generation methods. Each point is one patient. Gray represents real records from the SJLIFE cohort; blue, red and green represent simulated records using methods Random, Patterned and Frequent, respectively (see Methods). PC1, PC2 and PC4: first, second and fourth principal components.

## Conclusions

We introduced here a locality-sensitive hashing strategy for simplifying the analysis and comparison of semi-structured data, with an emphasis on electronic health records. The strategy involves reduction of potentially very complex data of arbitrary structure into small “fingerprints”, which are vectors of numbers of predetermined size. These vectors can then be easily compared in pairwise fashion or as a group using PCA or other ordination methods. While our novel methodology is currently designed for semi-structured data, it works well also for fully structured data, and (based on initial experiments) can be extended to work with fully unstructured data.

As one practical example, many healthcare questions can benefit from the ability rapidly detect similarities between records in sets with millions of patient records. The technique can be used on EHR data alone without domain knowledge, but, if needed, it could easily be combined with domain-specific pre- and post-processing to improve results. Recently, a different type of locality-sensitive hashing (minhash) was applied to detecting similarity between clinical notes [10]; our method supports analysis and comparison of complete electronic health records.

We presented here examples of analyses of electronic health records, but our approach is applicable to a multitude of different domains in which semi-structured data are commonly used.

## Methods

### Overview

Our algorithm identifies local units of structure in semi-structured data objects, expresses each such unit as a ‘triple’, described below, converts each triple into a vector of weights and collects these vector representations of triples into a ‘raw’ data fingerprint, which is then normalized. The resulting ‘normalized’ fingerprint can then be used to compare between data objects by simple correlation.

### Computation of data fingerprints

#### Recursion over JSON/XML structure

We use standard parsers to interpret arbitrarily complex JSON and XML input data into nested data structures composed of the basic data types: scalars (including numbers, strings, booleans and nulls), arrays (ordered lists of zero or more values, each of which can be of any type) and objects (unordered collections of key-value pairs, where the keys are strings and the values can be of any type). We then recursively study the resulting data structure to identify and enumerate patterns of local substructure, which we encode as ‘triples’ (see below). Each local structure pattern is independent of the rest of the structure, and therefore the order in which the overall structure is studied (width-first or depth-first) is not important. In practice, the order tends to be depth-first.

#### Flattening of uninformative structure layers

As a special case that arises frequently when analyzing XML objects, we ignore uninformative intermediary layers of structure that consist of an array with a single non-scalar entry. Thus, data structures matching the pattern *element_1_*->[0]->*element_2_* are treated in the same way as *element_1_*->*element_2_*.

#### Fingerprint length L

The output of the data fingerprinting method is a vector of numbers. The main parameter of the method is the length of this vector, which we denote *L*. It is possible (and frequently desirable) to compute in parallel fingerprints with different values of *L*, from the same input data.

#### Computation of vector values for the various data types

Vector value computation has three steps: 1) determine the type of the scalar value, 2) compute a raw vector value (the method depends on the scalar data type, as described below), and 3) normalize the raw vector value by stretching it to a minimal value of zero and a total weight of one.

#### Numbers

A number is represented by separately encoding its mantissa and its exponent. The (signed) mantissa of the number is multiplied by *L*/10 to be a number in the (-*L*..+*L*) range, then encoded into the vector value, modulo *L*. The (signed, integer) exponent is directly encoded into the vector value, modulo *L*.

#### Strings

Strings are encoded as a weighted Markov chain of the numeric values (native 8-bit encoding, like ASCII or Unicode) of each character in the string. We add to the vector value the numeric value of the first character in the string (modulo *L*) and use this character to initialize a growing string; the value of subsequent characters is weighted against that of the growing string with a user-settable weight defaulting to 0.1.

#### Encoding structures as triples

A core concept of the method is the transformation of JSON structures (objects and arrays) into collections of triples; a raw fingerprint is the sum of the vector values of all triples identified in the JSON structure. Each triple captures local structure and its associated values. triples are enumerated from JSON objects and arrays as follows. *Objects*. JSON objects are associative arrays: unordered collections of key-value pairs, equivalent to ‘hashes’ in Perl and ‘dictionaries’ in Python. Such objects are encoded into a fingerprint as the sum of the triples [objectName, key, value], where ‘objectName’ is the name of the object in the JSON structure. triples for which value is zero or null can be optionally skipped. The value of an object is the number of triples used.

#### Arrays

JSON arrays are most simply interpreted as ordered lists of elements, but depending on context could be interpreted as unordered lists or as sets. Since this information is not intrinsically represented in the JSON structure, arrays are by default interpreted as ordered lists, but can be optionally interpreted unordered lists or sets.

An ordered array A = [A_1_, A_2_, …, A_len_] is encoded into a fingerprint as the sum of the triples [*arrayName*, 0, A_1_], [*arrayName*, A_i_, A_i+1_] and [*arrayName*, A_*len*_, *len*], where *len* is the length of the array, i takes the values [1..*len*-1] and *arrayName* is the name of the array in the JSON structure. The value of an array is the number of triples contributed: 1+*len*.

An unordered set S of size *len* is encoded into a fingerprint as the sum of the triples [*setName*, 0, S_i_], where i takes the values [1..*len*] and *setName* is the name of the set in the JSON structure. triples for which value S_i_ is zero or null can be optionally skipped. The value of a set is the number of triples used.

#### Computing the vector value of a triple

The vector value of a triple [X, Y, Z] is computed as the average of [X, *σ*(Y), *σ*(*σ*(Z))], where *σ*(x) represents the left circular shift function.

#### Fingerprint normalization

Fingerprint vectors are normalized by subtracting their mean and dividing by their standard deviation.

### Fingerprint comparison

The methods for comparing data fingerprints are the same as described for genome fingerprints [11] and genotype fingerprints [12]. Pairwise comparison of two fingerprints of the same length is achieved by computing the Spearman correlation between the two vectors. We provide efficient software for comparing one query fingerprint to a database of fingerprints, and to perform all-against-all comparisons within a database. Fingerprints can be collected into a data matrix, and then analyzed using PCA, t-SNE and other ordination methods.

### Validation data

#### SyntheticMass data

We obtained a sample set of 130,833 simulated patient records from SyntheticMass [6] via https://syntheticmass.mitre.org. The records are in FHIR Draft Standard for Trial Use (DSTU) 3 format. We computed data fingerprints with vector length *L*=200 and excluded 14,648 patient records with fewer than 250 triples. We performed PCA using the R function call prcomp(M,center=TRUE,scale.=TRUE).

#### SJL500 data

We considered data on demographics, symptoms and treatment for 500 patients from the St Jude Lifetime Cohort (SJLIFE), represented in JSON format. We computed data fingerprints with vector length *L*=200 and performed PCA as above.

### Artificial data generation methods

We used three resampling strategies to simulate data preserving different attributes of a template data matrix consisting of records (rows) containing data fields (columns). An important aspect of the data was that data was missing from multiple columns in a non-random pattern across rows. Method 1, “Random”, simulated rows by resampling (with replacement) from each column of the template dataset. This preserves the frequency of missing values, but not the pattern of missing values. Method 2, “Patterned”, preserved patterns of missing data by selecting (with replacement) a template row, and resampling (with replacement) each value present in the template row from the non-missing values in the same column. Method 3, “Frequent”, preserved primarily the pattern of missing data alone by selecting (with replacement) a template row, and replacing every present value with the most common among the non-missing values in the same column.

### Availability

Software for computing data fingerprints is freely available through the project page: http://db.systemsbiology.net/gestalt/data_fingerprints/ and on GitHub: https://github.com/gglusman/data-fingerprints.

## Author contributions

GG conceived of the study. GG, MR designed and implemented the system. GG, MR, AK, JY, EWD, JH performed analyses. JH, JY, SH contributed expertise in clinical records. GG, JH, MR wrote the manuscript. All authors edited and approved its final version.

## Acknowledgements

This work was supported by NIH grant U54 EB020406 and by the National Center For Advancing Translational Sciences of the National Institutes of Health under Award Number OT3TR002026. The content is solely the responsibility of the authors and does not necessarily represent the official views of the National Institutes of Health.

The St. Jude Lifetime Cohort Study was supported by NIH grant U01 CA195547.

